# Subjective confidence modulates individual BOLD patterns of predictive processing

**DOI:** 10.1101/2024.02.23.581697

**Authors:** André Hechler, Floris P. de Lange, Valentin Riedl

## Abstract

Humans are adept at extracting and learning sequential patterns from sensory input. This ability enables predictions about future states, resulting in anticipation both on a behavioral and neural level. Stimuli deviating from predictions usually evoke higher neural and hemodynamic activity than predicted stimuli. This difference indicates increased surprise, or prediction error signaling in the context of predictive coding. However, interindividual differences in learning performance and uncertainty have rarely been taken into account. Under Bayesian formulations of cortical function, surprise should be strongest if a subject makes incorrect predictions with high confidence.

In the present study, we studied the impact of subjective confidence on imaging markers of predictive processing. Participants viewed visual object sequences of varying predictability over multiple days. After each day, we instructed them to complete partially presented sequences and to rate their confidence in the decision. During fMRI scanning, participants saw sequences that either confirmed predictions, deviated from them, or were random. We replicated findings of increased BOLD responses to surprising input in the ventral visual stream. In line with our hypothesis, response magnitude increased with the level of confidence after the training phase. Interestingly, the activity difference between predictable and random input also scaled with confidence: In the anterior cingulate, we found tentative evidence that predictable sequences elicited higher activity for low levels of confidence, but lower activity for high levels of confidence. In summary, we showed that confidence is a crucial moderator of the link between predictive processing and BOLD activity.

## Introduction

Our visual experience evolves as a continuous stream of sensory states. In the natural world, states close in time are related, providing a probabilistic mapping of visual input from one moment to the next. Studies presenting temporal sequences of visual stimuli show that humans and animals track the underlying transitional probabilities (Sherman et al., 2020; Turk-Browne et al., 2009). After a training phase, both behavior and neural patterns indicative of prediction (or anticipation) emerge: Participants detect and categorize expected stimuli faster (Turk-Browne et al., 2010) and stimulus templates of expected stimuli can be decoded from fMRI activity (Kok, Jehee, et al., 2012; Kok et al., 2014). Interestingly, the activity in visual cortices elicited by a given stimulus increases when its occurrence violates previously presented patterns (Kaposvari et al., 2018; Manahova et al., 2018; Meyer & Olson, 2011; Richter et al., 2018). This difference is thought to represent the extent of surprise – the mismatch between expected and actual observations. A more specific interpretation is prediction error (PE) signaling as proposed by predictive coding (Rao & Ballard, 1999). In this framework, PEs are feedforward signals carrying the difference between feedback predictions and sensory input (Keller & Mrsic-Flogel, 2018).

Most studies reporting corresponding BOLD or EEG activity focused on manipulating the transitional probabilities of the sensory stream. However, the human inference process is subject to uncertainty (Hasson, 2017; O’Reilly, 2013), possibly implemented as a Bayesian integration of prior knowledge and current evidence (Knill & Pouget, 2004; Pouget et al., 2013). Consequently, Bayesian predictive coding suggests that the magnitude of PEs depends both on the reliability of the evidence (the consistency of patterns and the signal-to-noise ratio) and the precision of our predictions (the probability of a prediction among all hypotheses) (Jiang & Rao, 2022 but see Aitchison & Lengyel, 2017 for non-Bayesian interpretations). The latter can either be inferred with computational modeling or by recording participants’ subjective confidence in a given decision.

Confidence ratings are usually tied to an overt or covert decision by the participant. For this reason, they are rarely assessed when studying the learning of temporal patterns, which often aims at implicit processes (Schapiro & Turk-Browne, 2015; Turk-Browne et al., 2010). Nevertheless, a promising line of studies combined both aspects by interjecting explicit predictions and confidence ratings between long blocks of visual stimulation. The authors found that these ratings reflect the confidence of an ideal Bayesian observer during the stimulation phase (Bounmy et al., 2023, Meyniel et al., 2015; Meyniel & Dehaene, 2017). This suggests that subjective ratings are a reliable indicator of the underlying inference process. Consequently, we used confidence ratings to quantify individual levels of uncertainty during sequence learning.

In the present study, we aimed to show that PE activity scales with subjective confidence, building on a previous study using visual streams of everyday objects (Richter et al., 2018). Furthermore, we addressed whether PE activity emerges as a function of increasing activity for surprising input (surprise enhancement) or decreasing activity for predictable input (expectation suppression). Evidence for the latter is inconclusive in humans with few fMRI studies looking into the alternative explanations (Feuerriegel et al., 2021). To address this gap, we included three experimental conditions regarding the visual stream: Fully predictable, fully unpredictable (random), and surprising. Following previous work, activity decreases for predictable compared to unpredictable stimulation served as evidence for expectation suppression (Manahova et al., 2018; Ramachandran et al., 2017).

## Results

### Confidence increases for predictable input and explains improvements in behavior

For the visual stimulation, we used sequences of visual images comprised of five everyday objects in full color. Prior to the scanning session, participants completed a three-day training phase with 20 minutes of stimulation per day (Figure 1). This phase only included predictable and unpredictable sequences. After each day, we sampled the learning process with a sequence completion task: Participants saw incomplete sequences and had to choose the correct trailing object from a set of five options. Each trial was followed by a confidence rating prompt. We found that average confidence regarding predictable sequences increased over the training phase while it remained constant for unpredictable ones (Figure 2a). Objective accuracy in completing predictable sequences also increased over days (Figure 2b) and was highly correlated with confidence (r=0.8, p<0.001). For the following analyses, we used the average confidence rating in predictable sequences after the last training day. To ensure visual fixation and attentiveness during the stimulation blocks, we instructed participants to react to upside-down images with a button press (the *cover task*). Neither reaction time nor accuracy were significantly different between conditions (reaction time: t(*41*)=-1.2, p=.24; accuracy: t(*41*)=1.55, p=.13).

**Figure 1.**
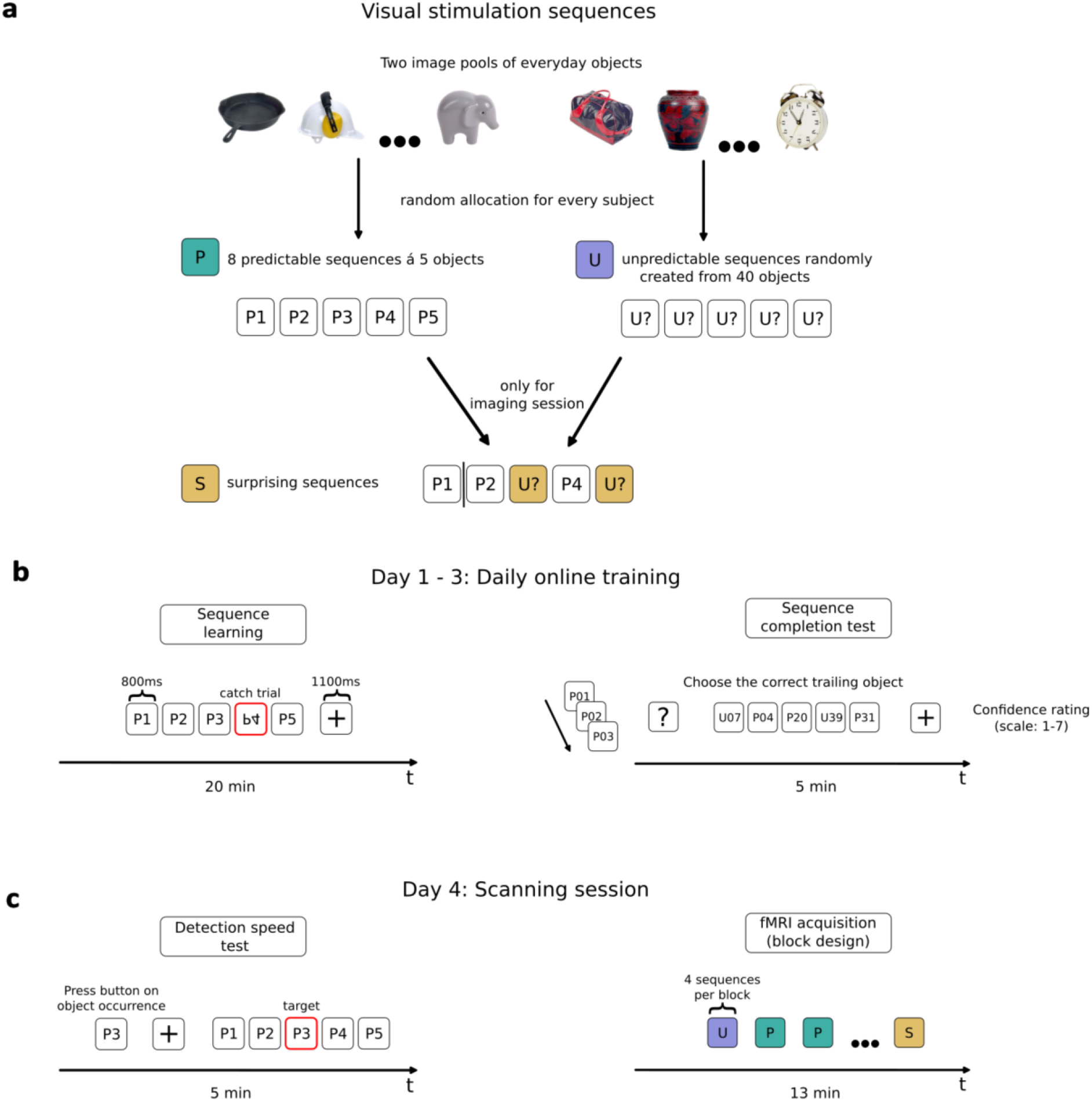
Experimental design. **a**. Creation of three types of object sequences. Predictable (P, green) and unpredictable (U, purple) sequences were shown during training. Surprising (S, yellow) sequences were only shown during the scanning session. See methods section for details. **b**. Across three days, participants performed an online training phase. Each day, sequences were shown for a total of 20 minutes. There was no inter-stimulus interval and sequences were separated by a fixation cross. To ensure gaze fixation and vigilance, participants were instructed to press a button when upside-down objects occurred (catch trials). During the following sequence-completion task, one to four of the leading objects were presented and the correct trailing object had to be selected from five options. The selection was followed by a confidence prompt. This task was not timed. Both predictable and unpredictable sequences were shown to keep object exposure comparable. **c**. Before entering the scanner, we tested for behavioral effects of sequence learning. Instead of reacting to upside-down images, participants had to detect and react to a target object which was given before each sequence. Both P and U were presented to quantify the gain in reaction time for predictable sequences. Finally, the scanning session followed the design of the learning phase with additional surprising sequences. We used a block design with four sequences per condition and a random condition order. There was no sequence completion test during this day.

**Figure 2.**
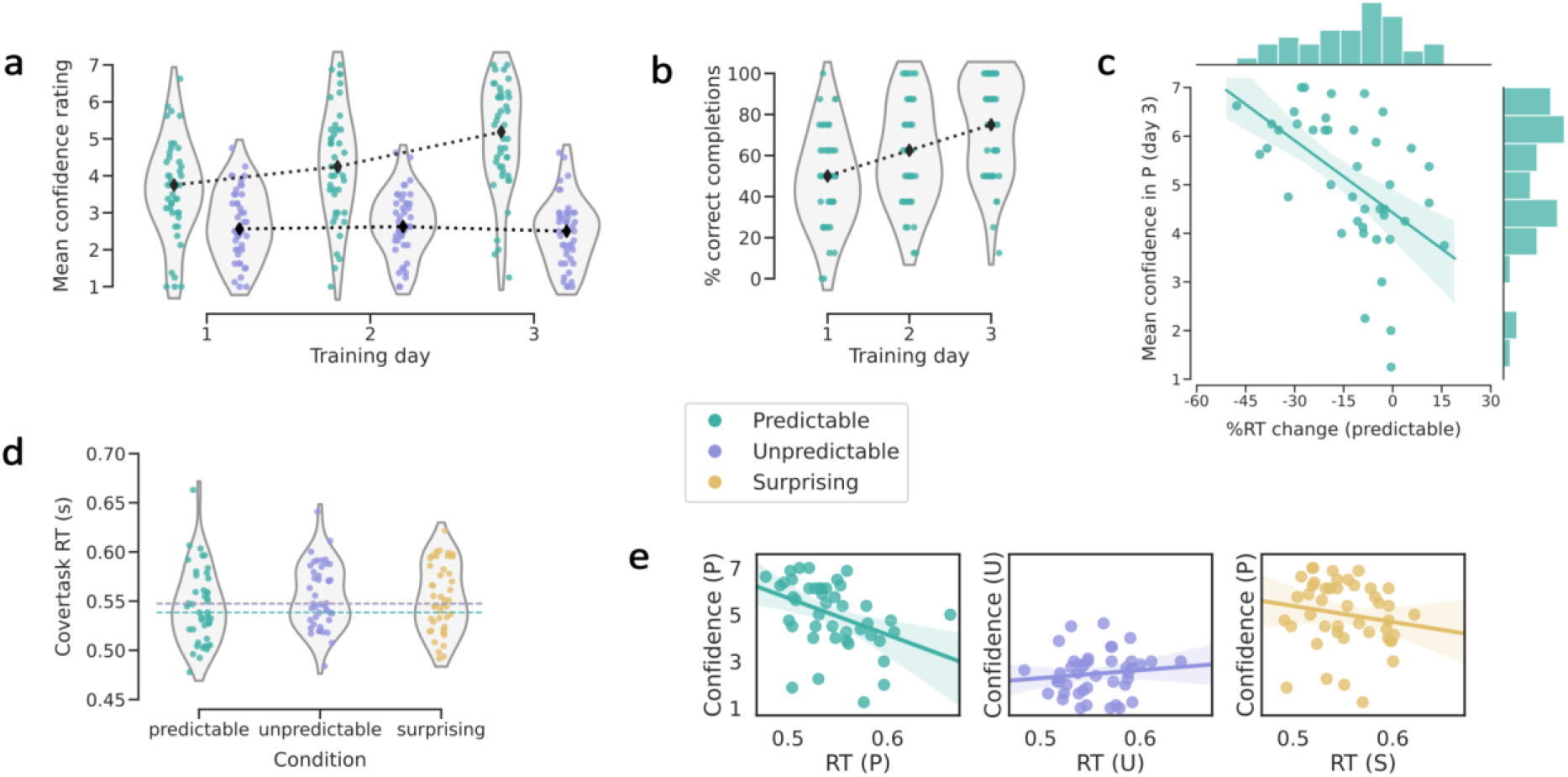
Training and cover task results. **a**. Average subject-wise confidence ratings within conditions after respective training days. Black markers show daily medians. **b**. Average percentage of correctly completed sequences. Unpredictable sequences could not be predicted over chance level and are not shown here. **c**. Significant linear regression between the relative difference in reacting to predictable versus unpredictable target objects and average confidence ratings for predictable sequences prior to scanning (r=-0.536, p<0.001). Negative values on the x-axis correspond to quicker reactions for predictable objects. **d**. Subject-wise averages of reaction time in the cover task during fMRI acquisition. Dashed horizontal lines show respective medians, with a significant difference between the predictable and unpredictable condition (post-hoc t-test: t(*41*)=2.59, p_FWE_=0.04). Note that the lines for the unpredictable and surprising conditions overlap since the median is nearly the same. **e**. Condition-wise correlations between confidence ratings and reaction time in the cover task during scanning. Since surprising sequences were based on predictable ones during scanning, we used the corresponding confidence ratings. The relationship was only significant for the predictable condition (r=-0.5, p_FWE_=0.002). The color legend applies to all subfigures.

We also tested whether learning the transitional probabilities led to behavioral advantages. To this end, we included a short task before the MR acquisition on day four. We presented the participants with a target object that would occur in the following object sequence. Sequence presentation followed the same design as in the training phase, but instead of performing the cover task, participants were instructed to react to the occurrence of the target object. Reaction times were nearly 10% faster for predictable objects (median RT change=-9.93%, t(41)=-5.66, p<0.001), with a maximum gain of 48%. Confidence was a crucial predictor of this effect: Subjects with higher ratings after training had significantly larger gains in reaction time (Figure 2c).

During fMRI scanning, we used a block design with surprising sequences in addition to the conditions from the training phase. These were based on predictable sequences from the learning phase but had one to three objects replaced (Figure 1a). The scanning session included no sequence-completion task or confidence ratings. Analysis of the cover task revealed that reaction times were faster in predictable than unpredictable blocks (Figure 2d). The same trend was present, but not significant, compared to the surprising blocks (t(*41*)=1.94, p_FWE_=0.18). The accuracy of reactions did not vary between conditions. Since the cover task was independent of the presented condition, we explored whether reaction times were differentially impacted by confidence levels. We calculated condition-wise correlations and found that high confidence in the predictable sequences was associated with quicker reaction times (Figure 2e). This link was not significant in the other conditions. Seeing this possibly confounding effect, we included reaction times in the GLMs of the following BOLD analyses.

### Prediction errors in sensory areas and prediction activity in parietal areas

We analyzed two contrasts with respect to the BOLD data: A PE contrast (surprising > predictable) and a prediction contrast (predictable > unpredictable). We adopt the terminology of “prediction contrast” in reference to non-error related aspects of predictive processing. Confidence was included as a regressor of interest and reaction time differences between the respective conditions as a confound regressor (Methods). First, we inspected the main contrasts between conditions. PE activity was present bilaterally throughout the ventral stream, in close correspondence to previous work using an event-based design after implicit statistical learning (Figure 3a, left; Richter et al., 2018). The prediction contrast did not provide evidence of expectation suppression, as we observed no significant negative clusters (i.e. predictable < unpredictable). Note that these results are based on a conservative whole-brain analysis. However, we found two positive clusters: A medial cluster in the posterior cingulate cortex and superior parietal cortex and a left lateral cluster covering parts of the superior parietal cortex, inferior parietal cortex and a small area of the dorsal stream (Figure 3b, left).

**Figure 3.**
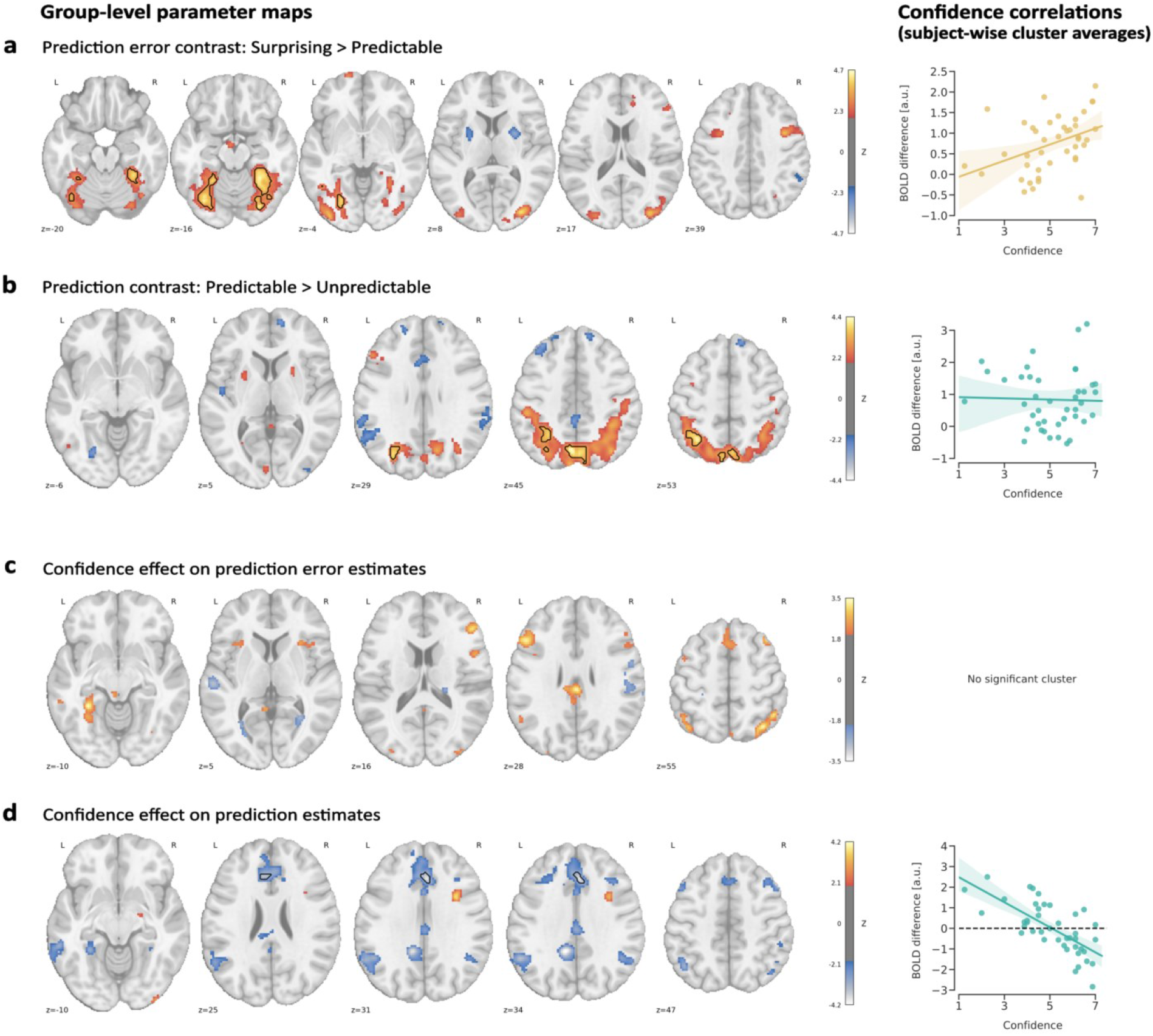
fMRI analysis results on the voxel level (brain slices, left) and cluster level (regression plots, right). Brain slices show uncorrected t-values thresholded at z=2.33 (p<0.01). Clusters surviving cluster correction (cluster-forming threshold: z=3.09, cluster threshold: p_FWE_=0.025) are shown in black contours. Regression plots show the subject-wise mean contrast parameter estimate across significant clusters and confidence ratings in predictable sequences after training. **a**. Surprising > predictable contrast. Confidence significantly explains interindividual differences in BOLD effects (R^2^_partial_ =.15, t(*41*)=2.54, p=.015). **b**. Corresponding results for the predictable > unpredictable contrast. The linear model yielded no significant results (R^2^_partial_ =.001, t(*41*)=-0.24, p=.82). **c** and **d**. Statistical maps for the confidence regressor in the respective GLMs. Brain maps indicate how strongly the BOLD differences between conditions vary with confidence levels. Cluster contours in **d** show a significant negative cluster (p=0.043) which did not survive FWE correction for two-sided cluster correction. Within this cluster, confidence explained considerable variance of the COPEs (R^2^_partial_ =.49). No test for significance was carried out because the ROI was based on the GLM results.

### Confidence explains interindividual differences in error and prediction activity

We investigated the link between confidence and predictive processing patterns in both a confirmatory and an exploratory way. Since we assumed that PE activity increases with confidence, we regressed the subject-wise average contrast parameter estimates (COPEs) in the significant PE clusters on confidence ratings, controlling for reaction time differences. In accordance with our assumptions, we found that PE activity significantly increases with the participants’ confidence level (Figure 3a, right). We then performed a corresponding analysis for the positive clusters of the prediction contrast. If these increases indicated PE activity, possibly due to uncertainty in the predictable sequences by low-confidence subjects, we would expect a similar relationship to confidence. However, we found no evidence for such a link (Figure 3b, right). A corresponding Bayesian correlation analysis revealed moderate evidence for the null hypothesis (BF=.195).

For the exploratory analysis, we analyzed the confidence regressor estimates of the second-level GLMs across the whole brain. No significant clusters emerged for the link between PE activity and confidence. The uncorrected maps indicate peaks in the ventral visual stream, anterior insula and inferior frontal cortex bilaterally (Figure 3c). Regarding the link to the prediction COPEs, we found a negative cluster in the anterior cingulate cortex (ACC; Figure 3d, left), although it did not survive FWE correction for two-sided cluster testing. Within this cluster, we found a strong negative association between COPEs and confidence. Interestingly, the corresponding regression line crosses the zero value with respect to the COPEs (Figure 3d, right). Descriptively, this indicates that the relative difference flips sign – for low-confidence participants, predictable blocks elicited higher BOLD signal compared to unpredictable blocks while the opposite happened for high-confidence subjects (Methods).

Finally, the directionality of the relationship with confidence generally depended on the contrast: Inspecting the uncorrected parameter maps, we see that PE activity increases with confidence while the inverse can be seen for the prediction contrast activity (with a single exception in the inferior frontal cortex, Figure 3d, left).

## Discussion

In the present paper, we established a link between brain activation patterns of predictive processing during the presentation of temporal sequences and subjective confidence in knowing the underlying patterns. Importantly, confidence ratings were derived separately from the visual stimulation. While this approach deviates from similar work using statistical learning, we replicated BOLD findings from a recent study where participants were kept unaware of the patterns (Richter et al., 2018). In line with our assumptions, PE activity in the ventral visual stream increased with confidence in knowing the (previously reliably predictable) object sequences. In the ACC, we found tentative evidence that the activity during predictable input relative to unpredictable input depended on confidence levels: For high confidence levels, activity for predictable input was lower. Conversely, for low confidence levels, unpredictable input elicited lower activity levels. These results stress the importance of accounting for interindividual differences in uncertainty during partly implicit predictive processes.

### Paradigms for learning of temporal regularities

The human ability to extract regularities in the visual domain has been studied from various perspectives, leading to subtly different paradigms (Conway, 2020). These include statistical learning (Fiser & Aslin, 2002), sequence or sequential learning (Gheysen & Fias, 2012), probabilistic learning (Meyniel et al., 2015) or temporal community learning (Schapiro et al., 2013). Instead of being fundamentally different, these approaches can be described as gradually varying along three axes representing the extent of exposure, structure, and instruction or feedback (Conway, 2020). Importantly, unifying frameworks have been developed, arguing that Bayesian inference models can account for most variants of acquiring regularities from sensory stimulation (Fiser & Lengyel, 2019; Konovalov & Krajbich, 2018). While the present study did not aim to resolve competing perspectives, it focused on the latter account. Here, we combined an extended learning phase of continuous sensory stimulation, using deterministic sequences and light instructions with a separate explicit performance task.

### Prediction error activity

We first discuss the results of our main contrasts with respect to previous studies. Firstly, we found evidence for PE activity throughout the ventral visual stream. Similar to previous studies in humans, there was no evidence of expectation suppression (Feuerriegel et al., 2021). In our data, the process behind PE activity is thus better explained as upregulated processing for surprising stimuli as opposed to downregulated processing for predictable ones. However, we here used a whole-brain approach that is more conservative than targeted analyses of the visual stream.

Our findings bear striking similarity to a previous study (Richter et al., 2018) despite differences in pattern length, instruction and participant awareness. This suggests that the underlying neural processes are not exclusive to incidental or implicit learning. Differences between neuroanatomical substrates of implicit and explicit learning point to more frontal activations for the latter, while implicit conditions predominantly affect sensory areas (Gheysen & Fias, 2012). However, these systems may work in parallel, with the implicit processes as a prerequisite for the explicit ones (Batterink et al., 2015). Consequently, one might expect additional activations in higher cognitive areas for our design. While not surviving cluster correction, we did find corresponding clusters in the precentral gyrus and inferior frontal cortex bilaterally (Figure 3a). It is possible that our study is underpowered to detect these effects. However, our GLM contrast was not designed to detect general patterns of statistical learning, but specifically isolated PE processing after a multi-day training phase. Related to the implicit versus explicit divide, there is evidence that surprising stimuli only evoke higher activity when unattended (Kok et al., 2012). In contrast, our data suggests that PE activity persists even when participants were explicitly asked to reproduce the patterns. This was corroborated by reaction time data in our cover task: Reaction times were shortest for predictable sequences, suggesting that, even though participants were highly engaged here, surprising sequences still evoked a stronger signal. Of note, our study differs from previous work by using a block design. That means that our surprising condition represents a visual stream including both confirmations and violations of expectation. The subtleties of transient, event-based responses might thus deviate from our findings.

### Prediction activity

Previous research on the contrast between predictable and unpredictable stimulation produced varied results. Stimuli predictive of other stimuli can elicit either increased activity (Egner et al., 2008) or decreased activity (Davis & Hasson, 2018), potentially dependent on task relevance (Richter & de Lange, 2019). Neuroanatomical findings partly overlap with the ACC being implicated irrespective of effect directionality. In our data, we found extended increases for predictable input focusing on the superior parietal cortex, intraparietal sulcus and posterior cingulate. The intraparietal sulcus has been found to increase activity in response to predictive stimuli (Egner et al., 2008) and is generally sensitive to the entropy of visual and auditory sequences (Nastase et al., 2014). Interestingly, the superior parietal sulcus is central to a wide range of predictive processing tasks, as shown by a recent modality- and task-general meta-study (Ficco et al., 2021).

### Interpreting the effect of confidence

Confidence ratings are often studied in the context of decisions, where they reflect the posterior probability of being correct, given evidence and an internal model (Pouget et al., 2016, but see Adler & Ma, 2018 for a non-Bayesian view). However, a line of studies (Bounmy et al., 2023; Meyniel et al., 2015; Meyniel & Dehaene, 2017) showed that confidence ratings can also be used to sample the uncertainty during continuous perceptual inference. This suggests that confidence is an ever-present aspect of perception, which can be cast as ongoing inference about the correspondence of our internal model (and the associated predictions) with sensory input (Friston, 2010). However, while many studies explored imaging patterns of expectation and surprise, surprising trials were often defined by the objective deviation of a presented stimulus from preceding patterns (Kaposvari et al., 2018; Manahova et al., 2018; Meyer & Olson, 2011; Richter et al., 2018). We suggest that much of the variance in individual response can be explained by accounting for individual (un)certainty in the presented patterns. As an example in the context of simpler paradigms, presenting a sequence of nine house images followed by a face image might lead to varying levels of surprise if observers did not infer the same conditional probability of observing a tenth house image. Given more complex patterns, as used in our study, interindividual variability is expected to increase. While the underlying reasons are beyond the scope of this study, the frequent divergence of human inference and behavior from (Bayes-) optimal models has been discussed previously (Acerbi et al., 2014; Beck et al., 2012).

Confirming our main hypothesis, PE activity in the significant clusters across the visual stream scaled with confidence. Previous studies investigating this link used model parameters derived from an ideal Bayesian observer, finding the strongest links between surprise and confidence outside of the visual cortex (Bounmy et al., 2023). To our knowledge, only two other studies explored the link between imaging indicators of PE and confidence ratings: In an EEG analysis of a perceptual decision task, parietal event-related potentials covaried with confidence ratings both during stimulus presentation and response (Boldt & Yeung, 2015). An fMRI study using a visual search paradigm found no evidence of PE scaling during stimulus presentation but did find an inverse relationship during the participants’ response in the inferior frontal gyrus (Sherman et al., 2016). To extend our findings, we performed an exploratory whole-brain analysis but did not find significant clusters. However, uncorrected results generally indicated an increase of PE activity with confidence across the cortex.

We also tested whether prediction activity scaled with confidence but found no relationship within the clusters of the contrast between predictable and unpredictable sequences. Mirroring the previous analysis, we then performed a whole-brain analysis on the voxel level. This yielded a cluster in the ACC where nearly 50 percent of the variance was explained by confidence levels. The direction of this effect was opposite to the PE contrast: Here, contrast estimates decreased as a function of confidence. Since this cluster did not survive correction for multiple comparisons, the results should be interpreted with care. However, evidence for the role of the ACC in predictive processing and conflict processing in general is widespread throughout the literature (Alexander & Brown, 2019; Garrison et al., 2013; Vanveen & Carter, 2002). Consequently, we discuss some implications of this finding in the following lines. Recently, the ACC has been described as part of a prefrontal network that anticipates PEs across cortical hierarchies (Alexander & Brown, 2018). This suggests a decrease in activity with increasing confidence, which has been confirmed for probabilistic learning tasks (Bounmy et al., 2023; Davis & Hasson, 2018). However, if the ACC is central to PE processing, we would expect corresponding results regarding our PE contrast. Possible explanations for this dissociation might lie in the hierarchical nature of predictive processing (Alexander & Brown, 2018): PEs in the visual stream indicate low-level mismatches without necessary awareness, while errors in higher cortices are subject to conscious reflection and, possibly, anticipation. Future studies are necessary to differentiate the modulating effect of confidence across the hierarchy of predictive processing.

Lastly, the confidence parameter estimates in the ACC flipped sign at medium-to-high levels of confidence. This means that predictable sequences elicited higher activity than unpredictable ones for low confidence levels, while the opposite was true for high levels. This matches the pattern usually expected for expectation suppression, albeit not in a sensory area. Regarding low levels of confidence, this can be explained by precision weighting of expected errors (Yon & Frith, 2021): If participants inferred that there is a ground truth to the predictable sequences while not being able to accurately predict the order, PEs of high precision can be expected. Due to the stochastic nature of the unpredictable condition, stimuli can never be accurately predicted, but the precision of errors is low. Nevertheless, if the unpredictable condition did elicit residual error activity, it might explain the relatively higher activity for high confidence levels. Here, the correspondence between predictions and sensory information is (close to) perfect in the predictable condition, triggering little to no error anticipation or processing. Finally, it is possible that even after three days of exposure, subjects still assumed that there is some order in randomness (Huettel et al., 2002), leading to incorrect predictions of comparatively high precision. Tentative evidence for this assumption can be found in the confidence ratings for the unpredictable condition. While these did not change during training, they were still far from minimal, indicating that some of the subjects did not correctly infer that these sequences were inherently unpredictable.

## Methods

### Participants

We recruited 44 participants including students, doctoral researchers, clinic staff and the general population of Munich. The sample size was based on a-priori analysis, aiming to detect at least a medium effect size (d>=0.5, alpha=0.05, beta=0.9). All participants took part in a familiarization MRI session (20 minutes), an online training phase over three days and the main MRI session (70 minutes) on the day after training completion. Two participants were excluded from the analysis due to technical problems during data acquisition. The remaining 42 participants (19 female, age [mean(std)] = 27.1(3.9)) were included for analysis. The study was approved by the ethics board of the Technical University of Munich (TUM), and we acquired written informed consent from all participants.

### Visual stimuli

We selected 224 full-color images of everyday objects from a larger image set (Brady et al., 2008). The stimuli were chosen to be maximally homogenous regarding salience, e.g., by excluding food items and bright colors. If not stated otherwise, all allocations of stimuli to subjects and conditions were random. A set of 80 images was selected for every subject, with half assigned to the predictable condition and the other half to the unpredictable condition.

### Experimental design

#### Implementation

We used Psychopy (Peirce et al., 2019) to implement the design. The online training sessions were realized using pavlovia.org, where Javascript-translated Psychopy experiments can be run online with millisecond precision (Bridges et al., 2020; Sauter et al., 2020).

#### Main task

Participants saw sequences of five objects each, each stimulus presented for 800ms with no inter-stimulus interval and a 1100ms fixation cross between sequences. Three types of sequences were presented: Predictable sequences had the same order for every repetition. Unpredictable sequences were composed of randomly chosen objects from the subject-specific set of unpredictable images. After all unpredictable images were shown, new sequences were formed. Surprising sequences were based on predictable sequences but had one to three stimuli replaced with images from the unpredictable condition for every repetition. To allow for an initial prediction, the first object was never replaced. The visual stream was reshuffled prior to presentation based on two rules: Objects could only repeat after all objects in the respective condition were presented, and objects (or sequences) could not appear twice in a row. This was done to minimize the confounding effects of repetition suppression. To ensure fixation and concentration, participants were instructed to quickly react to occurrences of upside-down objects. These appeared with a probability of 10%, independent of condition.

#### Training phase

Over the course of three days prior to the main scanning session, participants followed an online implementation of the main design. This phase only included predictable und unpredictable sequences. We instructed every participant to perform the training in a quiet environment without distractions. Stimulation blocks lasted 20 minutes per day, with a break of one minute after 10 minutes. Instructions were shown on screen before the stimulation began and the first day included a five-minute familiarization block with on-screen feedback regarding cover-task button presses. To measure objective and subjective progress in sequence learning, a testing phase followed each training day. Here, participants saw eight incomplete sequences for both conditions. After every sequence, participants chose what they assumed would be the correct trailing object from five options and gave a confidence rating on a scale of one to seven. No feedback was given regarding performance and no information on the underlying conditions was disclosed.

#### Imaging phase

During scanning, stimuli were presented against a grey background and subtended 4° of visual angle. We chose a block design with four sequences of one condition forming one block, with sequence order randomized. Half of the predictable and unpredictable objects were set aside for the generation of the surprising sequences. This ensured that every object was uniquely assigned to a condition. Participants saw 12 blocks per condition with a duration of 20.4s each, adding up to a run time of 12 minutes and 23.4 seconds. We disclosed no information regarding the sequence patterns and the imaging session included no sequence completion test.

### MRI acquisition

High-resolution structural scans were acquired with a T1-weighted 3D MPRAGE sequence (170 slices, voxel size = 1.0×1.0×1.0mm^3^, FOV = 250×256×170mm^3^, TR/TE/flip angle=9ms/4ms/8°). fMRI data was acquired using single-shot EPI (40 slices, voxel size = 3.0×3.0.3.0mm^3^, FOV 192×192×127.8mm^3^, TR/TE/flip angle=1200ms/30ms/70°) with 612 dynamic scans plus 2 dummy scans (total duration: 768 seconds).

### MRI data processing

We preprocessed the fMRI data with fMRIPrep (Esteban, 2019) which produces an automated processing description that we provide in the supplement.

### Statistical analysis

We performed whole-brain BOLD data analysis using FSL FEAT via a Python interface (adapted from Esteban et al. 2020). Regarding the first level analysis, we modelled the experimental manipulation as blocks, following their presentation timing with a length of 20.4 seconds (four object sequences) and included the respective temporal derivatives. The resulting boxcar functions were convolved with a double gamma hemodynamic response function. We added eight motion regressors: Six translations and rotations as well as two measures of bulk head motion (DVARS and framewise displacement). Finally, we included the average global CSF and white matter signal. All regressors were taken from the fMRIPrep results. The data were grand mean scaled, smoothed with a 6mm^3^ kernel and high pass filtered at 120 seconds. Since we did not have a resting condition, we created two GLMs: One leaving the surprising blocks unmodelled (used to create the prediction contrast between the modelled blocks) and one leaving the unpredictable blocks unmodelled (used to create the PE contrast). This was done to prevent overparametrization of the design matrix which can lead to unstable estimates.

Subsequent group-level analyses were run using FSL’s FLAMEO and included two covariates: Confidence ratings from predictable sequences (individual average after the last training day) and the percent change in reaction time to catch trials between the conditions being contrasted (individual average over all blocks). Since we acquired separate statistical maps from the first level analyses, we ran two corresponding analyses on the group level. This influences the interpretation of the covariates: The effect of confidence is now specific to the respective contrast (i.e. explained variance regarding the prediction contrast or PE contrast). Statistical maps of significant activation were obtained using FSLs cluster correction with a voxel threshold of z=3.09 and a cluster threshold of p=0.05 (Woo et al., 2014). Since cluster correction is applied for negative and positive clusters separately, we used an FWE-corrected threshold of 0.025. Our interpretation of contrast parameter estimates (COPEs) rests on the underlying subtraction of two parameter estimates. The resulting COPE thus informs both about the magnitude of the difference as well as the direction (which parameter weight is larger). Comparisons regarding the anatomical localization were based on relative voxel coverage over regions in the Harvard-Oxford Cortical Probabilistic Atlas.

All analyses on behavioral data were performed using the python package pingouin (Vallat, 2018). In linear regressions, the partial R^2^ values for confidence predictors were based on a partitioning of explained variance following Grömping (2006).

